# CLIMATE BRAIN - Questionnaires, Tasks and the Neuroimaging Dataset

**DOI:** 10.1101/2024.09.12.612592

**Authors:** Dominika Zaremba, Bartosz Kossowski, Marek Wypych, Katarzyna Jednoróg, Jarosław M. Michałowski, Christian A. Klöckner, Małgorzata Wierzba, Artur Marchewka

**Author notes:** corresponding author(s): Dominika Zaremba, Małgorzata Wierzba and Artur Marchewka.

## Abstract

Climate change presents a fundamental threat to human populations and ecosystems across the globe. Neuroscience researchers have recently started developing ways to advance research on this topic. However, validated questionnaires, experimental stimuli, and fMRI tasks are still needed. Here we describe the CLIMATE BRAIN dataset, a multimodal collection of questionnaire, behavioural, and neuroimaging data related to climate change, acquired from 160 healthy individuals. In particular, it includes data from (1) various questionnaire measures, including the Inventory of Climate Emotions (ICE); (2) a neuroimaging task for measuring emotional reactions to standardized Emotional Climate Change Stories (ECCS); and (3) a neuroimaging task based on Carbon Emission Task (CET) to measure climate action-taking. For technical validation, we provide image quality metrics and show the evidence for the effectiveness of tasks consistent with prior studies. To our knowledge, the proposed dataset is currently the only publicly available resource specifically designed to investigate human brain responses to climate change.

## Background & Summary

Climate change poses a significant and immediate threat to both human populations and ecosystems worldwide^1^. Despite clear expert recommendations, the pace of necessary systemic changes remains inadequate. Understanding human responses to climate change is crucial for fostering bottom-up, individual behaviour change, which is just as necessary as legal and technological solutions^2^.

Although many disciplines have contributed to this understanding, neuroscience can offer insights into the brain mechanisms that drive emotional reactions to climate change and pro-environmental decision-making. Research in environmental, affective, and decision neuroscience can deepen our understanding of the formation of pro-environmental attitudes and behaviour^3^. Integrating neuroscience and social sciences could lead to targeted interventions designed to promote sustainable behaviours and policies, ultimately contributing to global efforts to combat climate change.

One promising avenue of research involves the role of emotions related to climate change (henceforth: climate emotions) in the formation of attitudes and behaviours. Strong emotions, especially climate anger and climate hope have been the focus of researchers investigating drivers of environmental activism^4–6^, while climate anxiety has been identified as an important factor for youth mental health^7,8^. A better understanding of climate emotions can inform more effective strategies to motivate individuals and communities to engage in sustainable practices^9^. Affective neuroscience can provide another layer of explanation and bring new insights into the research and practice.

Many studies on environment and climate-related decision-making have traditionally relied on measures that, while cost-effective and convenient, often lack validity. Notably, conclusions about the effectiveness of interventions have been formulated based on self-report measures of behaviour, intentions, and attitudes^10^. Unfortunately, declarations and decisions made in hypothetical scenarios often poorly reflect real-life situations^11,12^. Thus, significant challenges persist in developing measures with higher ecological validity. A major step in this direction has been the development of the Carbon Emission Task^13^, a task that measures behaviour with tangible consequences. Here, we adapted this task to neuroimaging settings, providing a tool to investigate the neuronal mechanisms underlying climate-related decision-making.

Given that the efforts of environmental neuroscientists are pioneering, advancing this field requires adherence to open science practices, such as sharing validated research tools, stimuli and reliable data. Therefore, here we present the CLIMATE BRAIN dataset, a unique resource with climate-change-related data derived from 160 healthy participants.

CLIMATE BRAIN contains data from multiple sources, including the Inventory of Climate Emotions (ICE^14^) and other questionnaire measures, a neuroimaging task Reading and Rating Emotional Stories (RRES) designed to assess emotional responses to standardized Emotional Climate Change Stories (ECCS^15^), and the Carbon Emission Task (CET^13^), adapted to functional magnetic resonance imaging (fMRI) settings.

To the best of our knowledge, the proposed dataset is currently the only openly available source of data and tasks specifically designed to explore the relationship between emotion, pro-environmental decision-making, and other psychological, theoretically related individual characteristics, such as the level of climate change concern, sense of efficacy, socioeconomic status or political affiliation, on both behavioural and brain level.

## Method

### Subject characteristics

The study was conducted in accordance with the Declaration of Helsinki and was approved by the Ethical Review Board of the SWPS University in Poland (approval no. 2023-167). All participants provided written informed consent and signed an extended document detailing study information, including data privacy and the processes of pseudo-anonymization and anonymization for analyses and publications related to the research project. The consent form was inspired by the Open Brain Consent template^16^. The public dataset is now fully anonymized, ensuring the privacy and confidentiality of all participants.

The sample comprised 160 individuals aged 20 to 25 years, evenly split between men and women (80 each). Participants were recruited by an external company based on specific inclusion criteria: at least a high school education, no history of neurological or psychiatric disorders, right-handedness, no contraindications to MRI scanning, residence in Poland for most of the past five years and Polish as their first language. Additionally, they were required to exhibit a moderate level of concern about climate change, as indicated by their response on a scale from 0 to 4, where moderate concern was defined as a score of 1, 2, or 3. Participants received a fixed remuneration of 200 Polish zloty (PLN), approximately 46 EUR. However, in order to introduce experimental manipulation, participants were initially informed that they would receive a guaranteed 120 PLN, but their total remuneration could increase up to 240 PLN depending on their decisions in one of the tasks (see: Carbon Emission Task).

The gender, year of birth, place of residence, and education level of the participants were recorded. We also measured numerous variables that are theoretically related to climate emotion and decision-making measured in the current study, i.e. political affiliation, perceived socioeconomic status, driver’s license status, car usage frequency as a passenger or a driver, concern about climate change, climate action efficacy beliefs, perceived eco-friendliness of one’s lifestyle, perceived importance of individual or collective climate action, psychological distance to climate change, nature relatedness and baseline level of 8 climate emotions. Participants completed a health screening survey to ensure their safe participation in the experiment. The data from the health screening survey are not publicly available.

### Image acquisition

Neuroimaging data were acquired on a 3-Tesla Siemens Prisma scanner with a 32-receive channel head coil. An anatomical T1-weighted scan was acquired at the beginning of the scanning session using a magnetization-prepared rapid gradient-echo sequence (MPRAGE) with a voxel size of 1 × 1 × 1 mm isotropic (field of view = 256 × 176 × 256 mm [A-P; R-L; F-H]) in sagittal orientation. In the case of three participants, due to insufficient quality a T1-weighted scan had to be acquired for the second time at the end of the scanning session. Functional data were acquired using echo-planar imaging pulse sequence (multi-band acceleration factor 2, in-plane acceleration factor 2, repetition time [TR] = 2000 ms, echo time [TE] = 30 ms, flip angle [FA] = 70°). We acquired 66 slices in transverse plane orientation with an isotropic voxel size of 2.5 × 2.5 × 2.5 mm. For estimating magnetic field inhomogeneities, we additionally acquired two spin-echo EPIs with an inverted phase-encoding direction. The acquired DICOM files were converted to NIfTI and organized according to Brain Imaging Data Structure (BIDS) Specification version 1.8.035 using dcm2bids version 3.2.0^17^.

### Image processing

The released dataset comes from the preprocessing performed using fMRIPrep, 23.2.0a3^18,19^; RRID:SCR_016216, which is based on Nipype 1.8.6^20,21^, RRID:SCR_002502.

A B0-nonuniformity map was estimated based on two echo-planar imaging (EPI) references with topup^22^ in FSL^23^, version 6.0.7.7, RRID:SCR_002823. T1w images were corrected for intensity non-uniformity (INU) with N4BiasFieldCorrection^24^, distributed with ANTs^25^, version 2.5.1, RRID:SCR_004757 and used as T1w-reference throughout the workflow. The T1w-reference was then skull-stripped with a Nipype implementation of the antsBrainExtraction.sh workflow (from ANTs), using OASIS30ANTs as target template. Brain tissue segmentation of cerebrospinal fluid (CSF), white-matter (WM) and gray-matter (GM) was performed on the brain-extracted T1w using fast FSL. Volume-based spatial normalization to one standard space (MNI152NLin2009cAsym) was performed through nonlinear registration with antsRegistration (ANTs), using brain-extracted versions of both T1w reference and the T1w template. The following template was selected for spatial normalization and accessed with TemplateFlow^26^, 23.1.0,: ICBM 152 Nonlinear Asymmetrical template version 2009c^27^ [RRID:SCR_008796; TemplateFlow ID: MNI152NLin2009cAsym].

For each of the 4 BOLD runs found per subject (across all tasks and sessions), the following preprocessing was performed. First, a reference volume was generated, using a custom methodology of fMRIPrep, for use in head motion correction. Head-motion parameters with respect to the BOLD reference (transformation matrices, and six corresponding rotation and translation parameters) are estimated before any spatiotemporal filtering using mcflirt (FSL^28^). The estimated fieldmap was then aligned with rigid registration to the target EPI (echo-planar imaging) reference run. The field coefficients were mapped onto the reference EPI using the transform. The BOLD reference was then co-registered to the T1w reference using mri_coreg (FreeSurfer) followed by flirt (FSL^29^) with the boundary-based registration^30^ cost-function. Co-registration was configured with six degrees of freedom. Several confounding time series were calculated based on the preprocessed BOLD: framewise displacement (FD), DVARS and three region-wise global signals. The three global signals are extracted within the CSF, the WM, and the whole-brain masks. Additionally, a set of physiological regressors was extracted to allow for component-based noise correction (CompCor^31^). Frames that exceeded a threshold of 0.5 mm FD or 1.5 standardized DVARS were annotated as motion outliers. The preprocessed functional images were then smoothed with a 6-mm FWHM Gaussian kernel within SPM12 (Wellcome TrustCentre for Neuroimaging, University College, London, UK, http://www.fil.ion.ucl.ac.uk/spm/software/spm12) running on MATLAB2023a (MathWorks, http://www.mathworks.com).

### Dataset validation

Behavioural data analysis and visualisation were performed using R^32^ (Version 4.1.2) in RStudio^33^ (Version 2022.12.0, RRID: SCR_000432). Neuroimaging data quality measures were extracted with MRIQC software^34^. Neuroimaging data analysis was performed using SPM12 (Wellcome TrustCentre for Neuroimaging, University College, London, UK, http://www.fil.ion.ucl.ac.uk/spm/software/spm12) running on MATLAB2023a (MathWorks, http://www.mathworks.com). Additionally, the BIDS-Matlab^35^(RRID:SCR_022292) software was used to preprocess the confounds timeseries generated by fMRIprep. Finally, the neuroimaging results were visualized using the NIilearn^36^ (RRID:SCR_001362) toolbox for Python.

### Experimental Design

Before the MRI session, participants completed a demographic survey and a set of questionnaires. The MRI session began with the localizer, shim and T1-weighted scan sequences. After a brief instruction, they completed a training session followed by three target runs of **Reading and Rating Emotional Stories** (RRES) task. Then participants read detailed instructions for the **Carbon Emission Task** (CET) at their own pace. After answering comprehension questions and completing a training session, they completed the target CET run. In the end, a fieldmap scan was acquired. All instructions, training sessions and both experimental tasks were administered using Presentation software (Version 23.0, Neurobehavioral Systems, Inc., Berkeley, CA). The whole experimental session lasted approximately 1,5 hours, including a 1-hour-long MRI scan.

#### Questionnaires

Participants completed the Personal and Collective Action Efficacy^37^, Psychological Distance to Climate Change^38^, Nature Relatedness^39^, and Inventory of Climate Emotions^14^ questionnaires, and answered additional custom questions (see: Supplementary Materials) outside of the scanner.

#### Reading and Rating Emotional Stories (RRES)

The task was designed to measure brain activity related to experiencing climate emotions. We recorded brain activity while participants read short stories and rated their emotional response to them on the scales of valence and arousal. The stories were selected from the Emotional Climate Change Stories (ECCS) dataset^15^, which was previously found to be effective in evoking climate emotions.

Participants were pseudo-randomly assigned to three groups, each exposed to stories inducing either anger (ANG, n=54), hope (HOP, n=54), or a neutral state (NEU, n=52). Each participant read 12 target stories (ANG, HOP or NEU) and 12 control stories evoking a neutral state. In ANG and HOP groups, target stories were related to climate change and aimed to evoke anger and hope, respectively. In the NEU group, target stories described everyday situations without emotional content. Control stories used across all groups were identical, depicting everyday situations without emotional content. Stories were presented in three runs, with each comprising 8 stories (4 target, TAR and 4 control, CON) in a fixed order. After each story, participants were instructed to rate their emotions on two scales: valence, ranging from negative to positive emotions on an 11-point scale, and arousal, ranging from low to high arousal on another 11-point scale. They used a two-button response pad to move the cursor on the valence and arousal scales. An overview of the RRES is presented in Figure 1.

**Figure 1.**
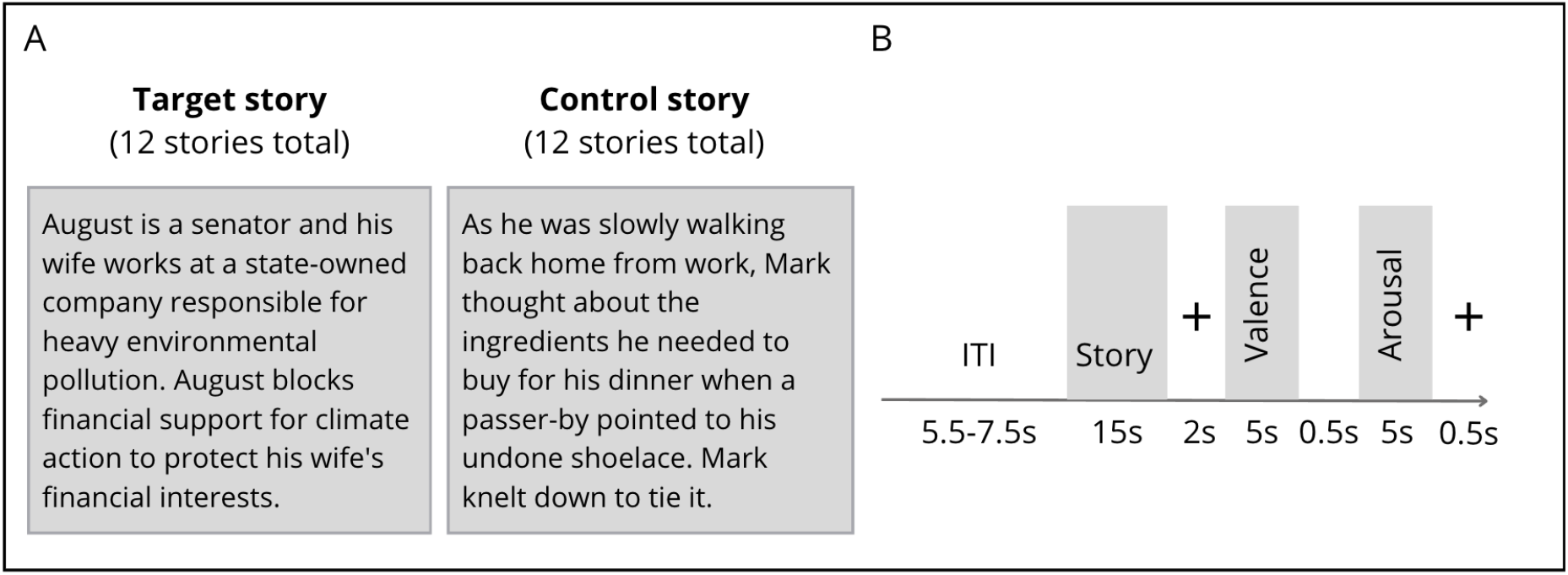
Reading and Rating Emotional Stories task overview. A. Participants read short stories about people and their actions. In ANG and HOP groups, target stories were related to climate change and aimed to evoke anger and hope, respectively. In the NEU group, target stories described everyday situations without emotional content. Here, we present an example of an anger-eliciting story and a control story. B. Stimuli were presented with jittered inter-trial intervals. Each trial consisted of two phases: the story-reading phase and the rating phase.

#### Carbon Emission Task (CET)

The Carbon Emission Task, developed by Berger and Wyss^13^, involves participants making a series of decisions between receiving a monetary bonus (a self-interest choice) and retiring carbon emission certificates (a climate-friendly choice). Choosing the climate-friendly option reduces the number of certificates available for purchase in Emission Trading Schemes. Emission Trading is an effective strategy for limiting the amount of carbon emissions^40^, which makes the CET task an ecologically valid way to measure pro-environmental behaviour related to climate change.

We adapted the original CET task for fMRI settings by: (1) reducing the visual complexity of the task; choosing stimuli presentation time and inter-trial intervals to maximize the signal-to-noise ratio; increasing the number of trials; (4) adjusting the possible monetary bonus and carbon emission levels; (5) adding dummy trials to control for motor activity (“Choose the option on the left/right”).

Data collection was carried out in an uninterrupted run consisting of 48 trials (36 target trials and 12 dummy trials). In the target trials, participants were instructed to select either a financially rewarding or climate-friendly option using a two-button response pad. Available options were different in each trial: monetary bonus of 0, 10, 50, 80, 100, or 120 Polish zloty (PLN; 1 PLN ∼ 0.25 EUR) versus a reduction of CO_2_ emissions by 0, 2, 10, 25, 40, or 50 kg CO_2_. In dummy trials, the value of options was 0 PLN and 0 kg of CO_2_. Participants’ task was to follow the instruction on the top of the screen and select the option on the left or the right (Figure 2). To ensure that participants treated each decision independently, they were told that one of their choices would be randomly selected to determine whether they receive a bonus or retire carbon emission certificates. In reality, all participants received a fixed bonus of 80 PLN on top of their base remuneration of 120 PLN. Additionally, we eliminated 25 kg of CO_2_ on behalf of each participant.

**Figure 2.**
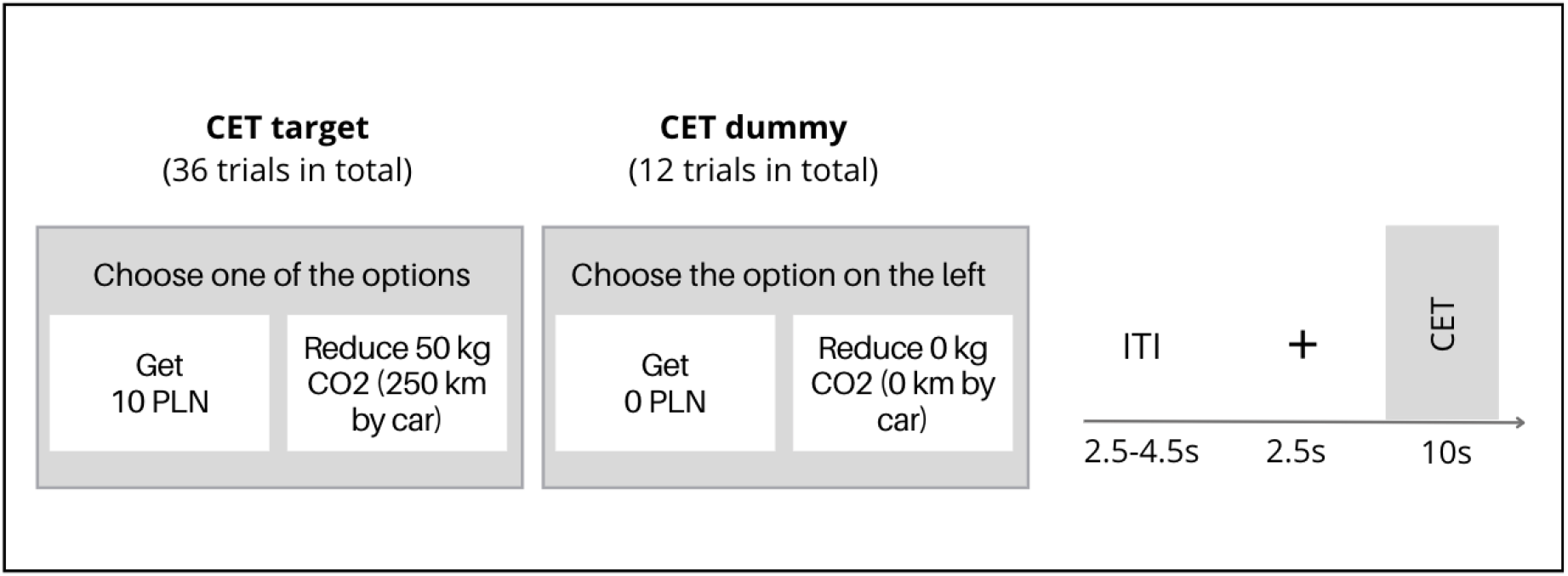
Carbon Emission Task overview. A. Participants made a series of decisions between a monetary bonus and a reduction of CO_2_ emissions. B. Stimuli were presented with jittered inter-trial intervals. Each trial began with a fixation cross, followed by a binary choice between a monetary bonus or carbon emission reduction.

### Data Records

This data descriptor outlines the behavioural and neuroimaging dataset available on OpenNeuro under the accession number ds005460 (https://openneuro.org/datasets/ds005460). The dataset is organized following the Brain Imaging Data Structure (BIDS) Specification version 1.8.035.

The `participants.tsv` file contains the demographic and questionnaire data of each participant. The accompanying `participants.json` sidecar file provides the metadata necessary to understand its content.

Additionally, the repository includes subject-wise `*_events.tsv` files, containing behavioural data for each task, along with corresponding `*_events.json` sidecar files that provide metadata necessary to understand their content.

The derivatives directory contains MRIQC and fMRIprep output. Due to anonymity protection, we do not share unprocessed brain images of the participants. In the case of MRIQC output, we share the summary file `group_bold.tsv`, which contains image quality metrics for all participants. In the case of fMRIprep output, we share subject-wise data organized in `sub-*` folders with preprocessed functional data, confound time series (allowing for motion and physiological noise correction), individual brain masks for each task and session, as well as accompanying `.json` sidecar files. These derivative files are named in the following convention:

- All files related to the RRES task contain the `task-stories` label in the filename.
- Additionally, files related to the respective sessions of the RRES task contain a run label: `run-01`, `run-02`, `run-03` in the filename.
- All files related to the CET task contain the `task-cet` label in the filename.

Supporting materials such as checklists used during experimental procedures, full demographic survey, questionnaires and task instructions are available in Supplementary Materials

### Technical Validation

#### Data collection and experiment design

The emotional stimuli come from a previously validated Emotional Climate Change Stories dataset ^15^. The Carbon Emission Task’s validity and reliability have also been already established ^13^. To ensure that the participants understood task instructions, we included a training session and comprehension questions in both RRES and CET tasks. To allow within- and between-group analyses, we included a control neutral group (NEU, n = 52) as well as control stimuli in each task. Personnel responsible for the data acquisition received detailed training and were required to check instruction comprehension during the data collection process (see Supplementary Materials).

#### Exclusion criteria

We used the MRIQC software to assess image quality, focusing on metrics such as temporal signal-to-noise ratio (tSNR) and framewise displacement. 4 participants were excluded from analyses due to excessive movement.

#### Behavioral and neuroimaging data

Participants from all groups exhibited similar climate change concern (Fig. 3) and had a similar profile of climate emotions (Fig. 4).

**Figure 3.**
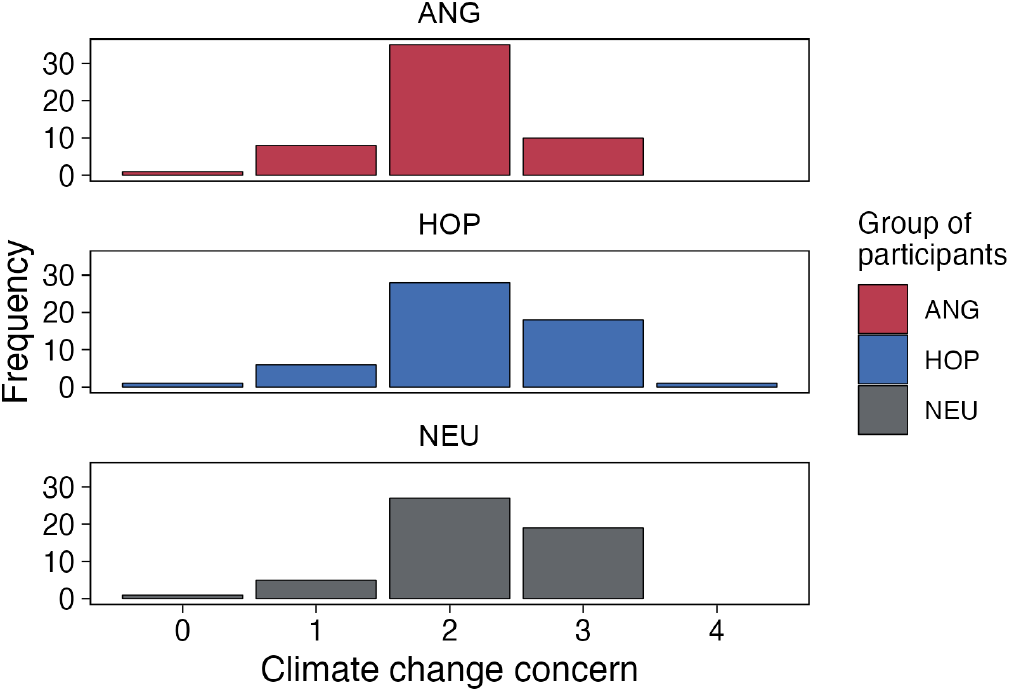
Climate change concern in each group (from no concern (0) to extreme concern (4)). No group differences were observed.

**Figure 4.**
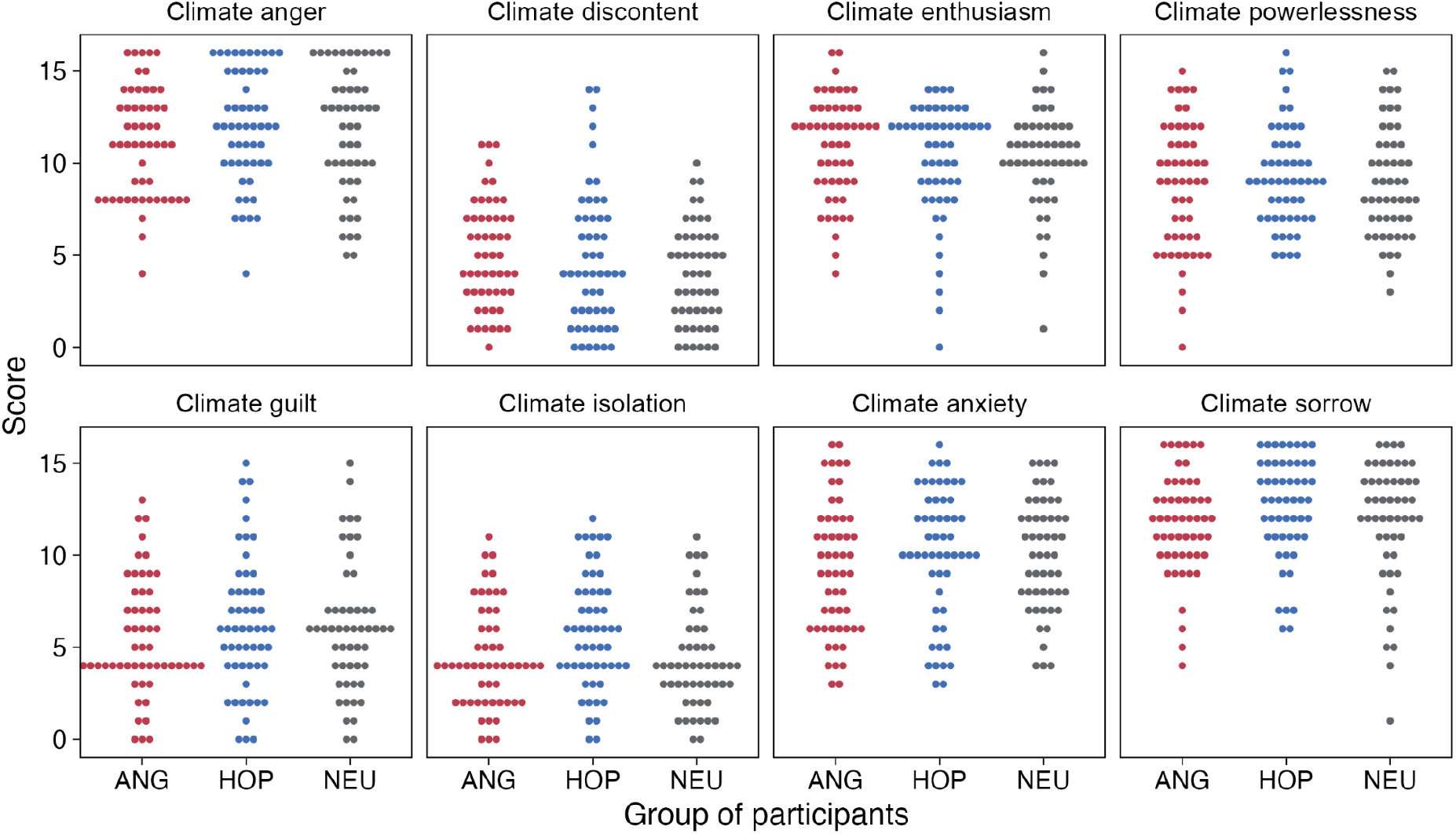
The profile of climate emotions assessed with the Inventory of Climate Emotions (ICE). Dots represent individual summary scores on each of the ICE subscales. Each subscale consists of 4 items, rated on a scale from 0-4.

### RRES validation

On the behavioural level, we checked the effectiveness of emotion elicitation in RRES by comparing the story ratings within and between groups. We have replicated previous results^15^ that the ECCS stories are effective in evoking emotional reactions that can be differentiated in terms of valence and arousal (Fig. 5).

**Figure 5.**
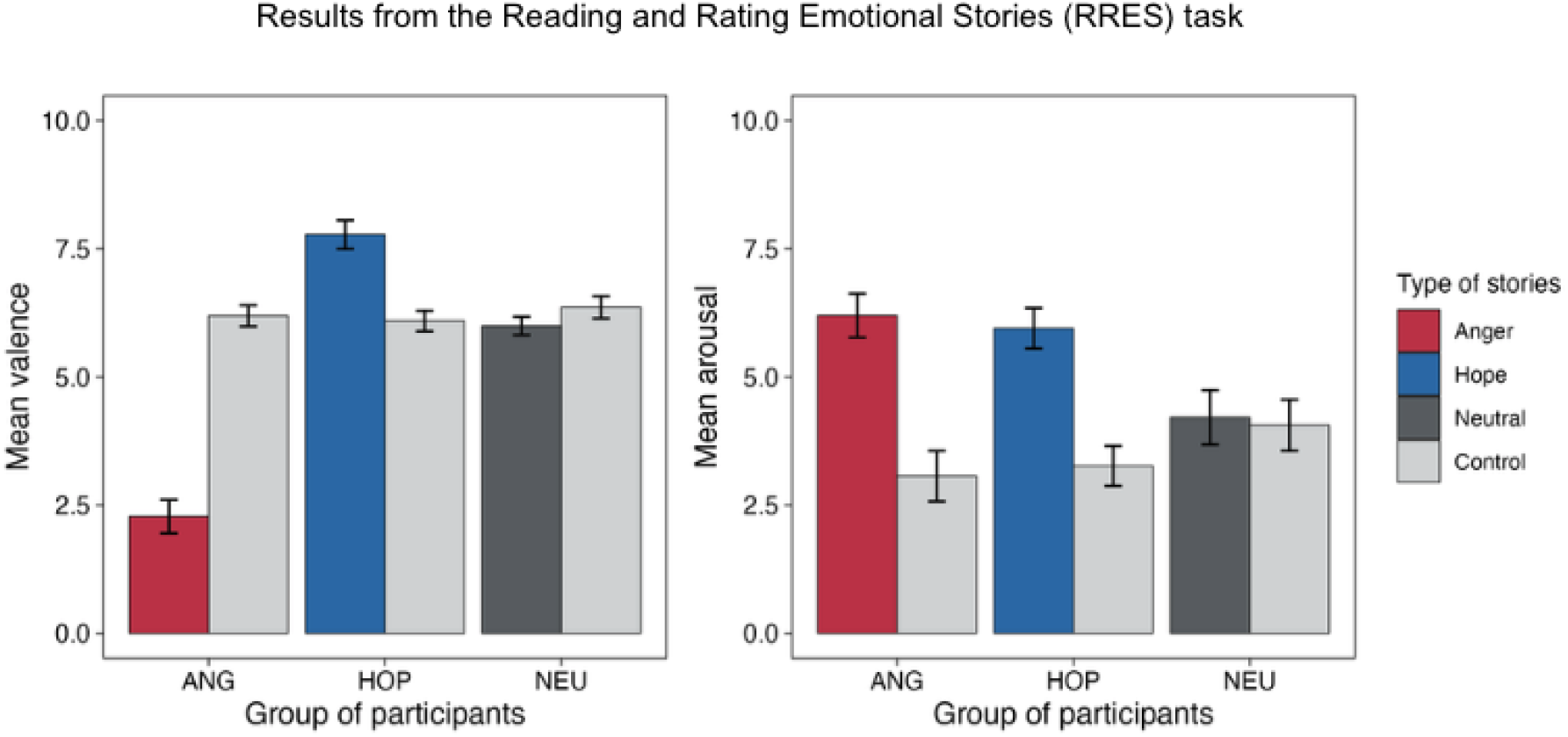
RRES validation. Mean arousal and valence ratings. Emotional stories are more arousing than neutral ones, anger stories are more negative, and hope stories - more positive. As expected, we do not observe differences between target and control stories in the NEU group. Note: Error bars represent 95% confidence intervals.

To validate the RRES task at the brain level we performed whole-brain analyses using a general linear model in SPM12. Timing corresponding to each event, along with multiple confounds were modelled at the subject level. The hemodynamic response was modelled with a canonical response function built into SPM12. Data were filtered with a 128 Hz high-pass filter. First-level contrasts comparing brain activation during reading target stories to a global baseline were entered into the second level one sample t-test models, separately for each group.

As expected, we observed increased involvement of brain areas within the reading network, including the ventral occipitotemporal cortex, inferior frontal gyrus, and superior and middle temporal gyri^41,42^. We also observed engagement of the limbic system, including brain areas such as the hippocampus, cingulate gyrus and insula^43,44^ (Fig. 6).

**Figure 6.**
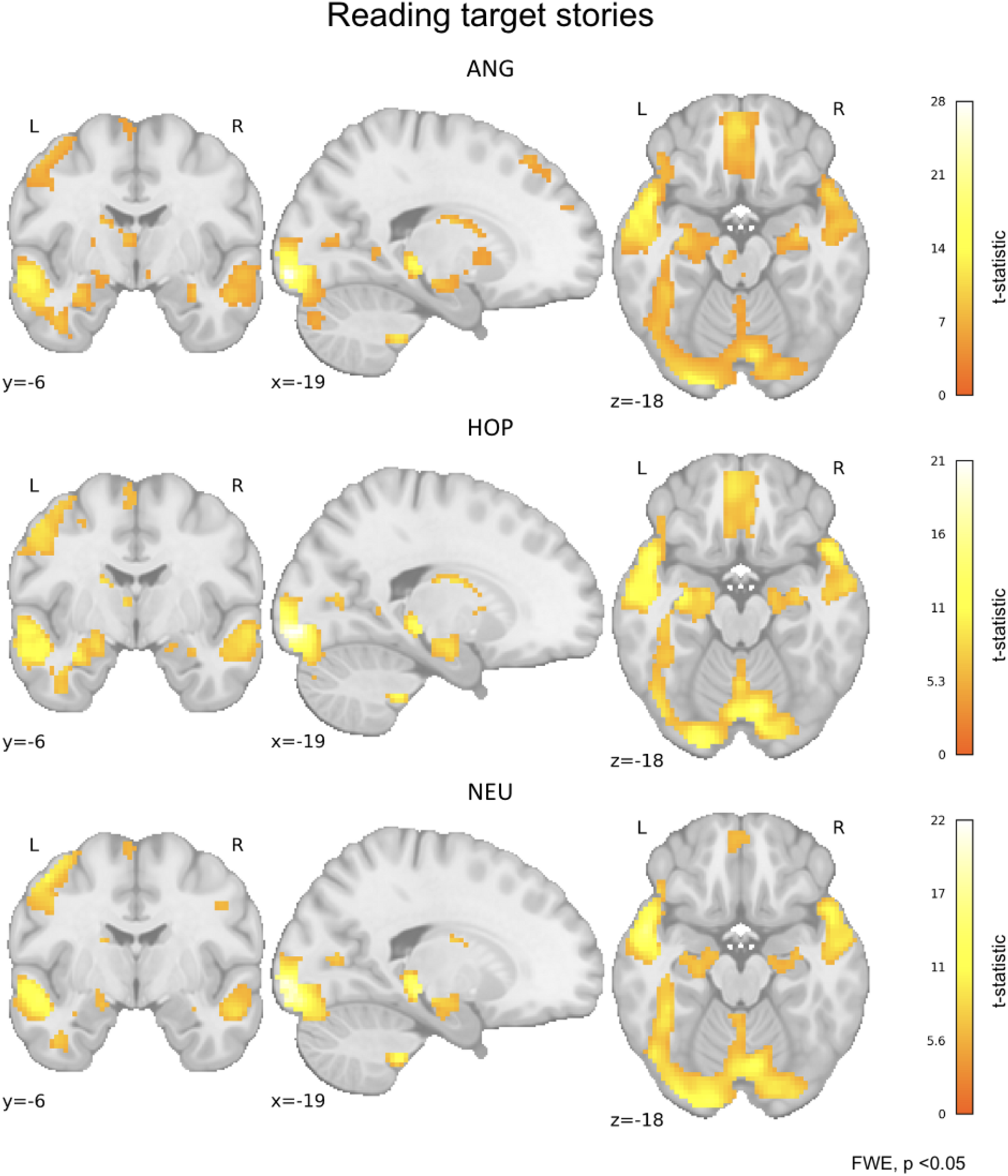
Regions of increased activity during reading target stories. Respective panels demonstrate the results for each group of participants: ANG (n = 54), HOP (n = 52), and NEU (n = 50). Statistical maps represent the contrasts for reading target stories (one sample t-test in each group). X, Y, Z—MNI coordinates. The colour bars represent t-statistic ranges. All maps are thresholded at p < 0.05, FWE (Family-Wise Error) corrected at the voxel level.

### CET Validation

To demonstrate the validity of the CET task, we conducted the behavioural analyses described in the original study^13^ and replicated previous results (Fig. 7). In particular, we showed that the proportion of climate-friendly choices linearly decreases with the size of the monetary bonus, and linearly increases with the amount of CO_2_ emission reduction.

**Figure 7.**
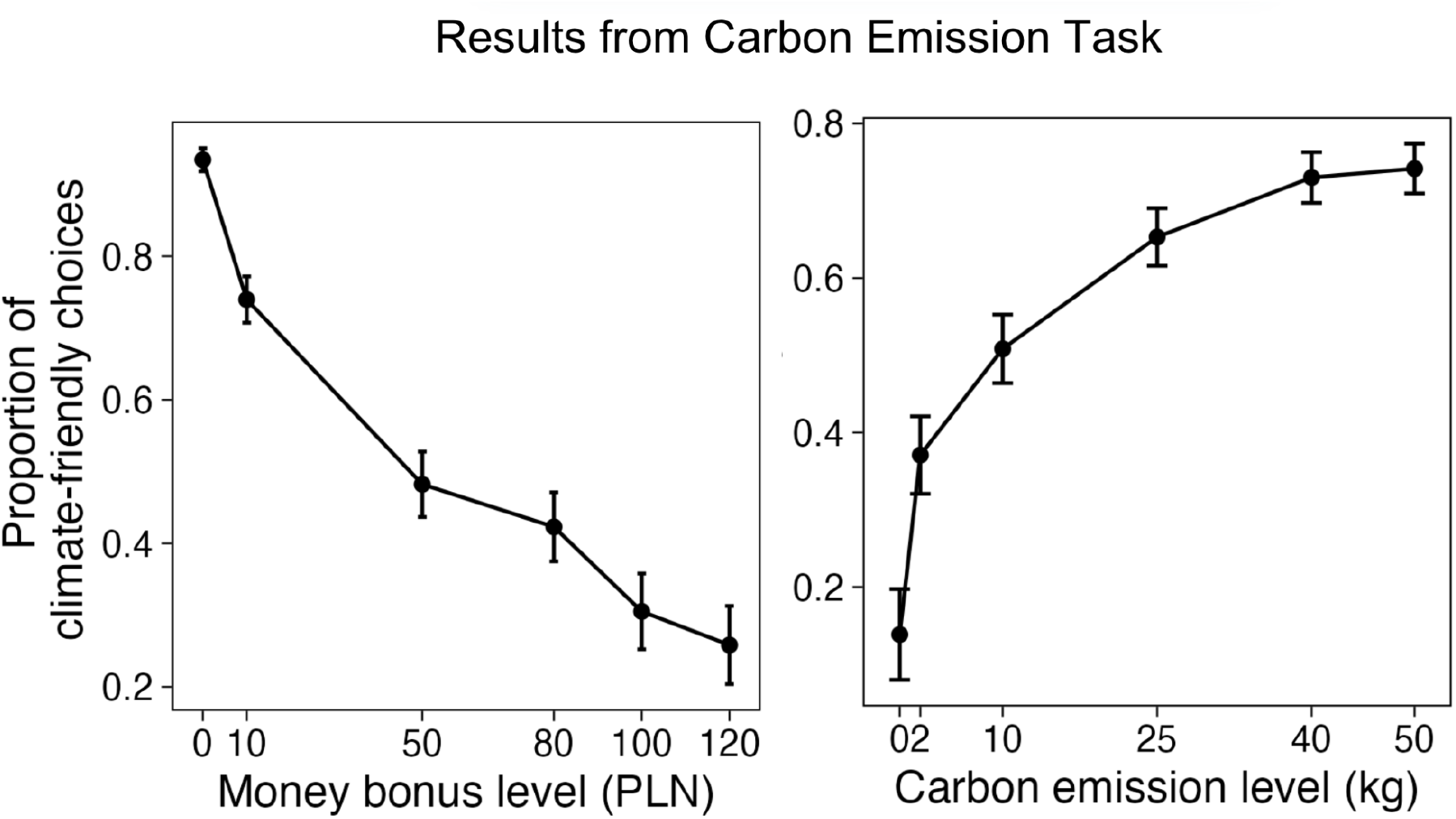
The proportion of climate-friendly choices depends on the size of the monetary bonus, as well as on how much CO_2_ emission can be reduced. Error bars indicate 95% confidence intervals.

To test the validity of the CET task at the brain level we performed a whole-brain analysis using a general linear model in SPM12. Timing corresponding to each event, along with multiple confounds were modelled at the subject level. The hemodynamic response was modelled with a canonical response function built into SPM12. Data were filtered with a 128 Hz high-pass filter. First-level contrasts comparing brain activation during climate-related decision-making target trials and dummy trials were entered into the second-level one-sample t-test model (n=156).

We observed increased activation in brain areas typically associated with decision-making, such as the superior and middle frontal gyrus, ventromedial prefrontal cortex, precuneus, hippocampus, posterior cingulate cortex, insula, and orbitofrontal cortex^45–47^ (Fig. 8).

**Figure 8.**
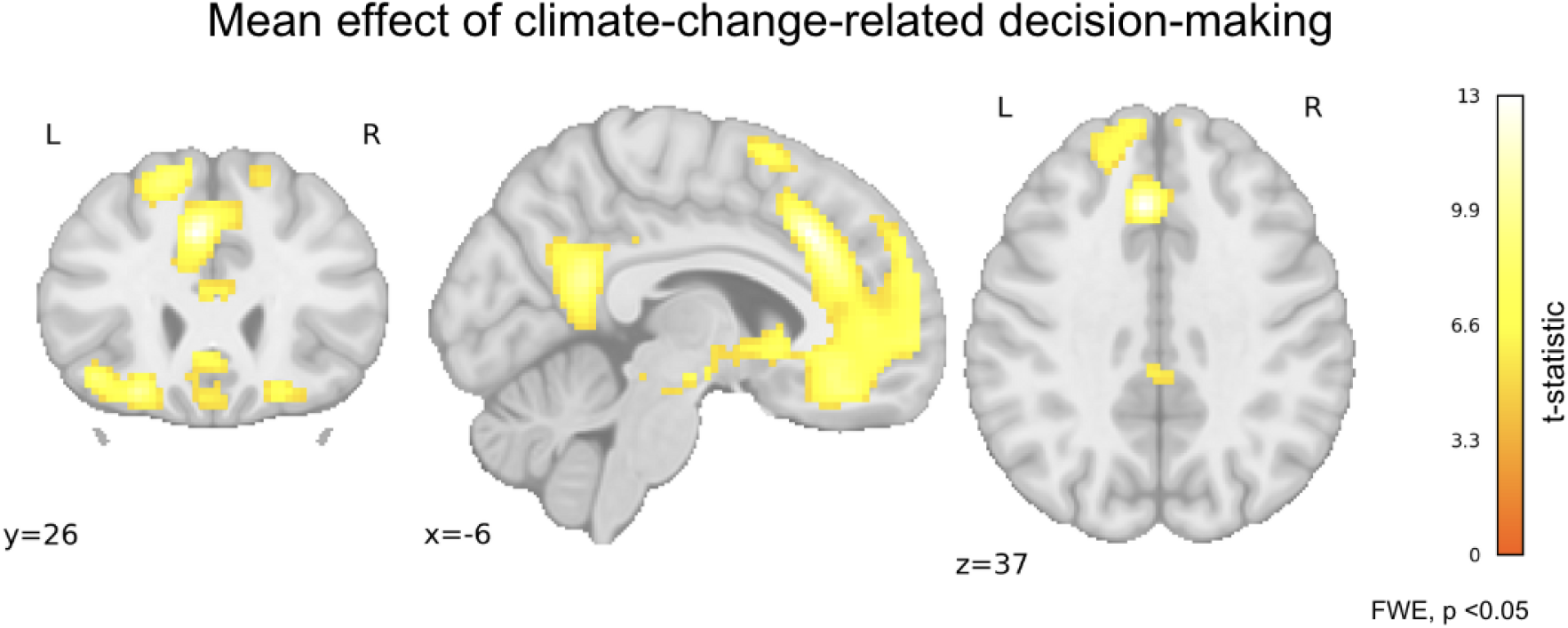
Regions of increased activity in climate-change-related decision-making. Statistical maps represent the contrasts for target trials compared to dummy trials (one sample t-test). X, Y, Z—MNI coordinates. The colour bar represents the t-statistic range. All maps are thresholded at p < 0.05, FWE (Family-Wise Error) corrected at the voxel level.

## Code Availability

All analysis code is available at https://github.com/nencki-lobi/climate-brain.

## Acknowledgements

We acknowledge the contributions of Michalina Marczak, the project initiator, Anna Czartoszewska, who supported the pilot experiments and as well as the research team who supported the data collection: Monika Tutaj, Anna Czartoszewska, Robert Szymański, Klaudia Chwiejczak, Weronika Kaczmarczyk, and Alicja Cieśla. We also extend our gratitude to all study participants.

The research conducted in this project is financed by Norwegian Financial Mechanism for 2014-2020 Grant No. 2019/34/H/HS6/00677.

## Author contributions

D. Zaremba: Conceptualization, Data curation, Formal analysis, Investigation, Methodology, Project administration, Software, Validation, Visualisation, Writing—original draft; Writing—Review and editing;

B. Kossowski: Conceptualization, Data curation, Formal analysis, Methodology, Resources, Software, Supervision, Validation, Writing—Review and editing.

M. Wypych: Methodology, Formal analysis, Writing—Review and editing;

K. Jednoróg: Conceptualization, Methodology, Writing—Review and editing;

J. M. Michałowski: Funding acquisition, Project administration, Writing—Review and editing;

C. A. Klöckner: Funding acquisition, Project administration, Writing—Review and editing;

M. Wierzba: Conceptualization, Data curation, Formal analysis, Investigation, Methodology, Resources, Project administration, Software, Supervision, Validation, Visualization, Writing—Review and editing.

A. Marchewka: Conceptualization, Formal analysis, Funding acquisition, Methodology, Project administration, Resources, Supervision, Writing—original draft; Writing—Review and editing;

