## Supplementary materials for "CLIMATE BRAIN - Questionnaires, Tasks and the Neuroimaging Dataset"

**RRES Instruction**

In a moment, you will read several stories describing various situations. All stories are based on real events and were created from the accounts of people who agreed to share their experiences with us.

Your task will be to read each story and then assess the intensity of your own emotions using the scales described below:

1. Do the situations described evoke rather negative or positive emotions in you?
2. To what extent do the situations described emotionally arouse you? Do you feel a complete lack of arousal, or strong arousal, agitation, or excitement?

Pay particular attention to the emotions you feel while reading the stories.

You will provide your answers using two buttons located on a special device. The left button will be used to change the answer to the left, and the right button to change it to the right.

You will always have the same amount of time to respond: 5 seconds for each question. After this time, your answer will be recorded, and you will move on to the next screen.

When responding, try to use the full range of the rating scale. There are no right or wrong answers; respond according to your first impression. You will start with a short training session.

**RRES Guide for the Experimenter**

**Initial Voice Instructions**
I will now display the instructions for the first task. You can scroll through them using any button.

**FAQ**

**What do the scales represent?**

On the first scale, you will rate whether your emotions are generally positive or negative, regardless of their intensity. On the second scale, you will rate how aroused or emotionally reactive you feel, indicating the strength of your emotional response.

**CET Instruction**

Carbon dioxide ($CO_2$) is considered a key factor contributing to climate change. Scientists around the world agree that climate change can only be mitigated if $CO_2$ emissions are significantly reduced.

To limit $CO_2$ emissions, a market for trading $CO_2$ emission allowances was created. Factories or power plants must purchase allowances, called emission certificates if they want the right to emit $CO_2$. The number of these certificates is limited.

For this study, we have purchased a certain number of $CO_2$ emission certificates from the market. Some of these certificates will be destroyed (permanently removed from the market). How many certificates are destroyed will depend on your decisions in the task.

If a $CO_2$ emission certificate is destroyed, no company will be able to emit that amount of $CO_2$. This will result in less $CO_2$ being released into the atmosphere, contributing to climate change mitigation.

In this task, you will be asked to make 48 decisions that could either reduce $CO_2$ emissions or increase your compensation.

To help you understand the consequences of your choices, we will additionally represent $CO_2$ emissions as the number of kilometres driven by an average car.

In each of the 48 trials, you will be asked to choose one of two options. You can either take a financial bonus ranging from 0 to 120 PLN or remove a certificate allowing the emission of 0 to 50 kg of $CO_2$.

In each trial, you will have 10 seconds to make a decision. If you do not decide within 10 seconds, the screen will automatically move to the next trial.

If you do not make a decision, you will not receive any money, and no $CO_2$ will be removed. Not making a decision is disadvantageous for both you and the climate.

Your choices do not accumulate. We will randomly select one of your decisions, based on which you will either receive a bonus or a $CO_2$ emission certificate will be destroyed. Depending on your decisions and the result of the draw: You could receive up to 120 PLN or remove up to 50 kg of $CO_2$, equivalent to driving 250 km by car.

Choosing to remove $CO_2$ increases the chances of destroying a $CO_2$ emission certificate, which helps mitigate climate change. Choosing a financial bonus increases your chances of receiving a payout. To check if you are paying attention to the task, sometimes we will ask you to choose a specific option, “Choose the option on the left” or “Choose the option on the right.” This instruction will appear at the top of the screen. You will not receive a bonus for these decisions, and they will not involve any $CO_2$ emissions. These decisions will not be included in the draw.

You can select the option on the left using the left button, and the option on the right using the right button.

**CET Guide for the Experimenter**

**Initial Voice Instructions**

This task includes detailed instructions. You can navigate through them by pressing the left and right buttons.
Once you reach the final screen, I will ask if everything is clear and address any questions you may have.

**Comprehension Questions**

1. **What will you be choosing between?**
   Accepted answers: money/remuneration/benefit for me and carbon dioxide/emissions/certificates/benefit for climate/environment
2. **How will your final remuneration be determined?**
   Accepted answers: it is not cumulative/they don't sum up + it is chosen randomly/it is drawn from all of my choices
3. **What should you do when you see a choice between 0 PLN and 0 kg CO2?**
   Accepted answers: follow the instructions at the top, choose left or right depending on the instruction
4. **What happens if you don't choose any of the options within 10 seconds?**
   Accepted answers: neither I nor the climate benefits/I don't receive money, and I don't reduce CO2 + the program shows the next choice

**FAQ**

1. **How much does a single CO2 certificate cost?**

I cannot provide such information at this time. Please focus on making your decision based on the information available about the amount of the premium, the kilograms of CO2 and the kilometres driven in the car.

**Tutorial corrective messages**

1. **Wrong choice in the dummy trial.**
   Please pay attention to the instruction at the top of the screen—if it says 'choose the left option,' select the option on the left.
2. **Clicking multiple times during the same trial.**
   Did you try to change your answer? Unfortunately, only the first response counts, so please make your decision carefully.
3. **No decision within 10 seconds.**
   If you don't choose an option, that decision will still be included in the final draw, and neither you nor the climate will benefit.
4. **Irrational choice in a trial c0_m2 (selects carbon reduction)**
   What was behind your choice in this trial? See, you could get a chance to get 50 PLN, but there was no risk that you will cause CO2 emissions. Is that clear to you?

**Debriefing**

We would like to inform you that in the last task, we had to tell you that the amount of compensation and emissions would be determined randomly to ensure the success of our experiment. However, in reality, we will make sure that neither you nor the climate will suffer any loss.

You will receive compensation of 80 PLN, making a total of 200 PLN for participating in the entire study. Additionally, on your behalf, we will destroy emission certificates equivalent to 25 kg of CO2, which corresponds to the emissions from driving 125 km by car.

**Full demographic and psychometric questionnaire**

**Demographic survey**

---

1. Please indicate your gender:

- Woman
- Man
- Other (Please specify: _______)

---

2. Please enter your year of birth:

- (Input field)

---

3. Which of the following best describes the area you live in?

- Large city
- Suburbs or outskirts of a large city
- Medium or small city
- Village
- Single farm or house in a rural area

---

4. What is your highest level of education?

- Incomplete primary
- Primary
- Lower secondary (Gimnazjum)
- Vocational or basic vocational
- Secondary
- Bachelor’s degree or engineering diploma
- Master’s degree or medical degree
- Doctorate, post-doctoral degree, or professor title
- Other (Please specify: _______)

---

5. How concerned are you about climate change?

- Not concerned at all
- Not very concerned

- Somewhat concerned
- Very concerned
- Extremely concerned

---

6. In your opinion, which actions have the greatest impact in combating climate change?

- Collective actions (governments, institutions, corporations) [1]
- Individual actions (ordinary people) [5]

*(Scale from 1 to 5 where 1 represents collective actions and 5 represents individual actions)*

---

7. Compared to others, how would you rate the environmental impact of your lifestyle?

- Definitely harmful to the climate [1]
- Definitely climate-friendly [5]

*(Scale from 1 to 5 where 1 represents harmful and 5 represents climate-friendly)*

---

8. Do you have a driver's license?

- Yes
- No

---

9. How often do you use a car (e.g., as a driver or passenger)?

- Once a year or several times a year
- Once a month or several times a month
- Once a week or several times a week
- Daily

---

10. Do you consider yourself as having specific political views?

- No
- Yes
    - If yes: Sometimes in politics, people refer to "left" and "right." Can you describe your views using these terms?
        - No
        - Yes

---

11. How would you describe your political views?

*(Visible only if the previous question's answers are both "Yes")*

- Left [0]
- Right [10]

*(Scale from 0 to 10 where 0 represents the left and 10 represents the right)*

---

12. Which of the following best describes your feelings about your current household income?

- We/I live comfortably on the current income level
- We/I manage on the current income level
- We/I struggle to manage on the current income level
- We/I find it almost impossible to manage on the current income level

---

**Perceived Climate Action Efficacy (PCAE)**

---

Instructions: Please indicate your level of agreement with the following statements.

---

Response Options:

- 1 - Strongly disagree
- 2
- 3
- 4
- 5 - Strongly agree

| |
|---|
| I believe my actions can have a beneficial influence on climate change. |
| Actions I take personally can help reduce the impacts of climate change. |
| Climate change can be averted by mobilizing collective effort. |
| If we act collectively, we will be able to minimize the consequences of climate change. |

**Psychological Distance (PD) to climate change**

---

Preamble:
Please indicate to what extent you agree or disagree with the following statements. Use the scale provided below.

---

Response Options:

- 1 - Strongly disagree
- 2
- 3
- 4
- 5
- 6
- 7 - Strongly agree

| |
|---|
| My local area will be influenced by climate change. |
| It will be a long time before the consequences of climate change are felt. |

**Nature Relatedness (NR)**

---

Title: What is Your Relationship with Nature?

---

Preamble:
Please rate the extent to which you agree or disagree with each of the following statements using a scale from 1 to 5. When responding, consider your true feelings rather than the feelings of "most people."

---

Response Options:

- 1 - Strongly disagree
- 2 - Somewhat disagree
- 3 - Neither agree nor disagree
- 4 - Somewhat agree
- 5 - Strongly agree

| |
|---|
| My ideal vacation spot would be a remote, wilderness area. |
| I always think about how my actions affect the environment. |
| My connection to nature and the environment is a part of my spirituality. |
| I take notice of wildlife wherever I am. |
| My relationship to nature is an important part of who I am. |
| I feel very connected to all living things and the earth. |

**Inventory of Climate Emotions (ICE)**

---

Preamble:
Perhaps you have heard that the Earth is currently experiencing climate change. The aim of this questionnaire is to examine your feelings on this subject.

---

Instructions:
Rate the extent to which the following statements apply to you. For each statement, select one answer on the scale from "strongly disagree" to "strongly agree."

This questionnaire is not intended to verify your knowledge, so there are no right or wrong answers. Choose the answer that best describes what you feel.

---

Response Options:

- Strongly disagree
- Somewhat disagree
- Neither agree nor disagree
- Somewhat agree
- Strongly agree

| | |
|---|---|
| **ANG14** | I feel angry that the political and economic system that we live in harms the climate. |
| **ANG13** | I am outraged that politicians allowed climate change to come this far. |
| **ANG10** | I feel outraged at corporations that harm the climate. |
| **ANG3** | I feel anger when I think of politicians who delay efforts to mitigate climate change. |
| **DIS5** | It annoys me to watch people succumb to climate hysteria. |
| **DIS7** | I am annoyed by the constant publicity around climate change. |
| **IND2** | I am bored of hearing about climate change. |
| **IND13** | I am surprised that people experience strong emotions in connection with climate change. |

| | |
|---|---|
| **EMP12** | The increasing public engagement with climate change gives me hope. |
| **HOPF9** | I believe that there are emerging solutions that will allow us to stop climate change. |
| **CHECK1** | To convince us that you are reading this, please, just mark the option “Strongly disagree”. |
| **HOPF8** | Concrete actions for the climate allow me to be optimistic about the future. |
| **EMP7** | Social mobilisation in the fight against climate change makes me feel that together we can achieve this goal. |
| **POWL11** | I feel confused about what I can do to reduce climate change. |
| **POWL7** | I am overwhelmed by how many aspects of life would need to be changed to limit climate change. |
| **POWL2** | As an individual, I feel powerless with little agency over what happens with the climate. |
| **POWL13** | I feel helpless when I think of how difficult it is to live in a climate-friendly way. |
| **GUI11** | I have a guilty conscience about not doing enough to mitigate climate change. |
| **CHECK2** | To convince us that you are reading this, please, just mark the option “Neither agree, nor disagree”. |
| **GUI6** | It upsets me that I have a big negative impact on the climate. |
| **GUI8** | I feel guilty that my lifestyle contributes to climate change. |
| **GUI12** | I am angry at myself for not doing enough to limit my negative impact on the climate. |
| **ISO4** | I feel like one of the few people who actually understand what climate change entails. |
| **CHECK3** | To convince us that you are reading this, please, just mark the option “Strongly agree”. |
| **ISO5** | I feel lonely because most of the people around me don’t care about climate change as much as I do. |
| **ISO8** | I feel lonely because it’s difficult to talk about my climate change concerns with other people. |
| **ISO12** | I feel alienated because society considers concern for climate change as something strange. |
| **APP7** | Thinking about climate change makes me fear for the future of our children. |

| | |
|---|---|
| **HOPL5** | I am overwhelmed by the awareness of the approaching climate disaster. |
| **HOPL11** | Everything seems uncertain because of climate change. |
| **APP14** | I fear how climate change will affect me and my loved ones. |
| **SOR13** | The thought of so many species going extinct under the pressure of climate change fills me with sorrow. |
| **SOR6** | The thought that the world I know is disappearing forever because of climate change makes me sad. |
| **SOR4** | I feel sorry about the possibilities we are losing forever because of climate change. |
| **SOR14** | I am sad that so many living creatures suffer because of climate change. |
